## Supplementary material for "GJA1 Depletion Causes Ciliary Defects by Affecting Rab11 Trafficking to the Ciliary Base"

**Supplements**

**Supplementary figure 1**

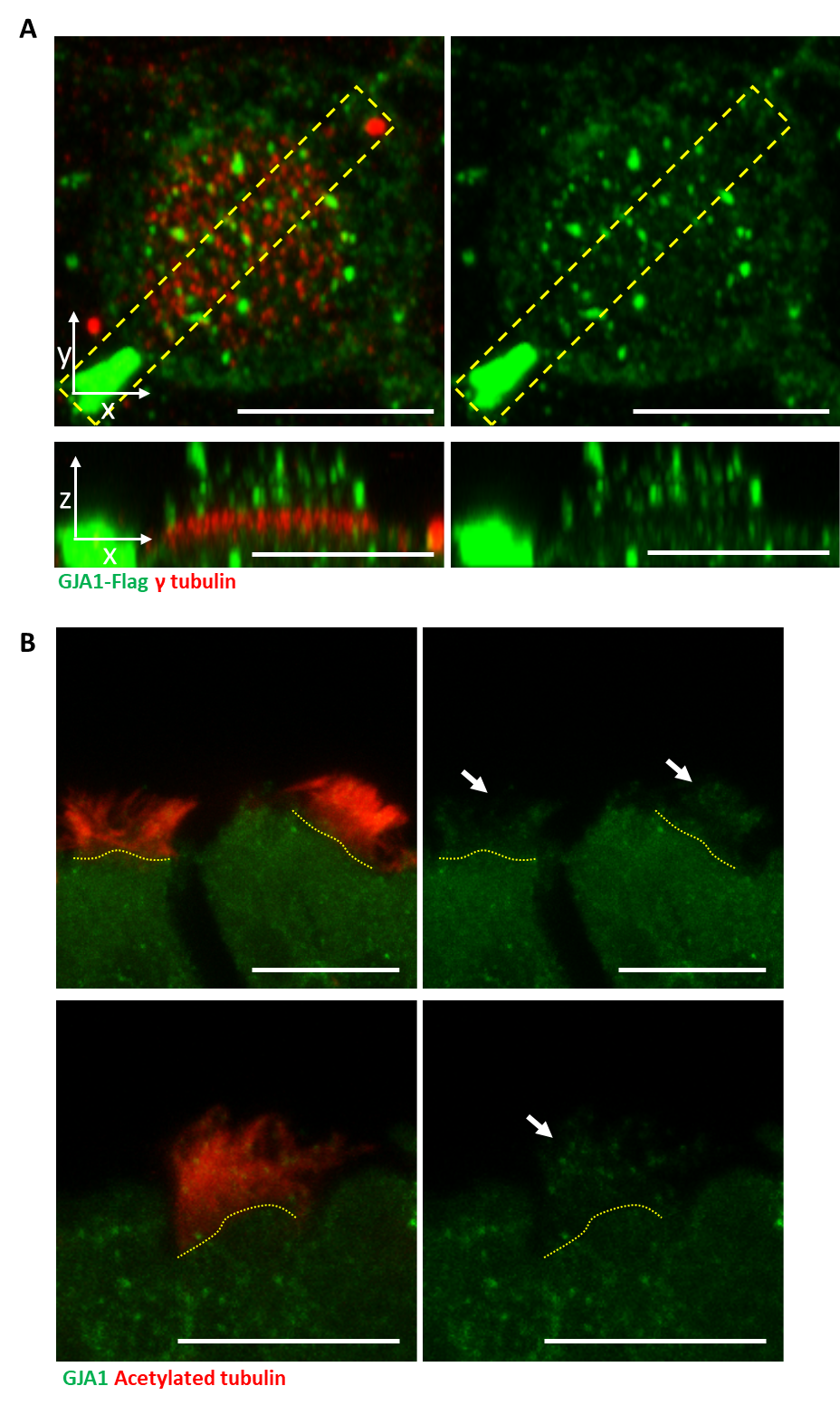

**Figure S1. GJA1 is localized to ciliary axonemes in *Xenopus* multiciliated cells and mouse tracheal tissues.**

**A.** Immunofluorescence analysis of GJA1 localization in *Xenopus laevis* multiciliated epithelial cells. *Xenopus laevis* embryos were microinjected with GJA1-Flag mRNA. The embryos were stained with antibodies against Flag (green) and γ-tubulin (red). The bottom images are X-Z projections of the top images. Scale bars: 10 µm.

**B.** Immunofluorescence analysis of GJA1 localization in mouse tracheal tissues. Mouse tracheal tissues were sectioned at 100 µm and stained with antibodies against GJA1 (green) and acetylated tubulin (red). Scale bars: 10 µm.

**Supplementary figure 2**

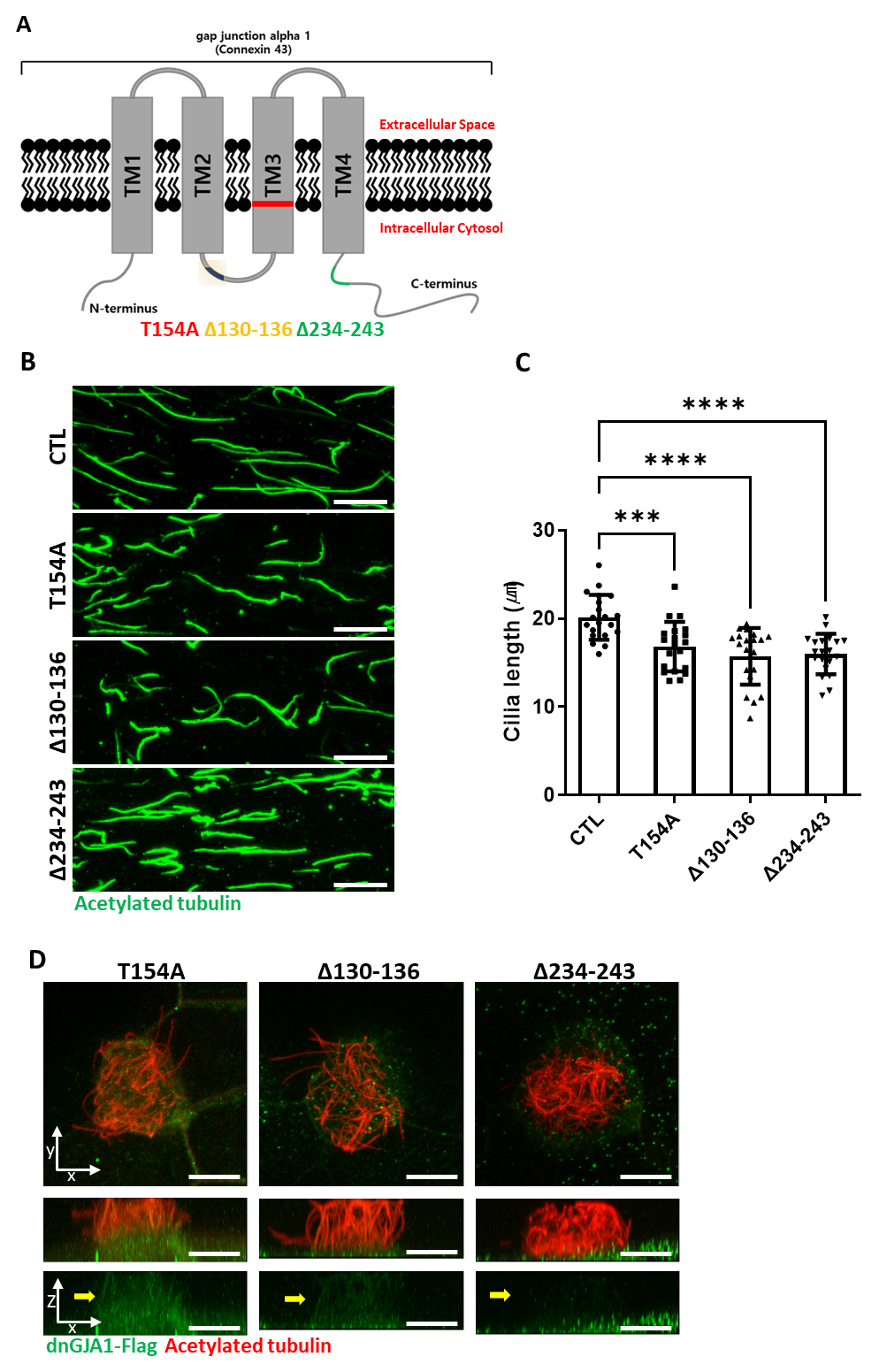

**Figure S2. Dominant-negative GJA1 mutants cause ciliary defects and exhibit differential localization.**

**A.** The schematic structure of dominant-negative GJA1 mutants (dnGJA1). The T154A mutant contained a threonine (154)-to-alanine point mutation in transmembrane domain (TM) 3. The Δ130–136 mutant contained a deletion of the intracellular loop between TM 2 and 3. The Δ234–243 mutant also contained a deletion of the C-terminus of GJA1.

**B.** dnGJA1-injected embryos were treated with cilia isolation buffer, and isolated cilia were labeled with an acetylated tubulin antibody (green). Scale bars: 10 µm.

**C.** The average lengths of ciliary axonemes in panel **(B)** were plotted. Error bars represent the mean±s.d. P values were determined by one-way ANOVA (P^***^=0.0008, P^****^<0.0001). Raw values are provided in the source data file.

**D.** Immunofluorescence analysis of the localization of each dnGJA1 mutant in *Xenopus laevis* multiciliated epithelial cells. *Xenopus laevis* embryos were microinjected with each dnGJA1-Flag mRNA, and the embryos were stained with antibodies against Flag (green) and acetylated tubulin (red). The bottom images are X-Z projections of the top images. Scale bars: 10 µm.

**Supplementary figure 3**

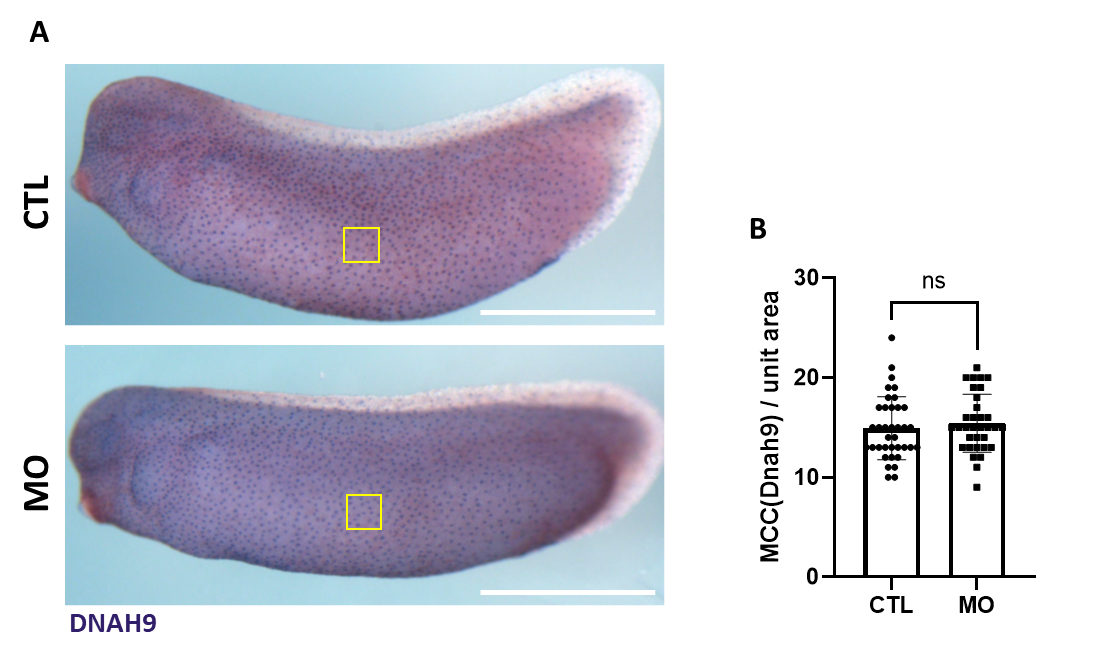

**Figure S3. MO-mediated knockdown of GJA1 does not affect the cell fate specification of multiciliated cells.**

**A**. Expression of a multiciliated cell fate marker, DNAH9, was not altered in GJA1 morphants. Scale bars: 1 mm.

**B**. Statistical analysis of DNAH9-positive cells in panel **(A)**. GJA1 depletion resulted in no significant difference in DNAH9 expression. P values were determined with a two-tailed t-test (P^ns^>0.05). Raw values are provided in the source data file.

**Supplementary figure 4**

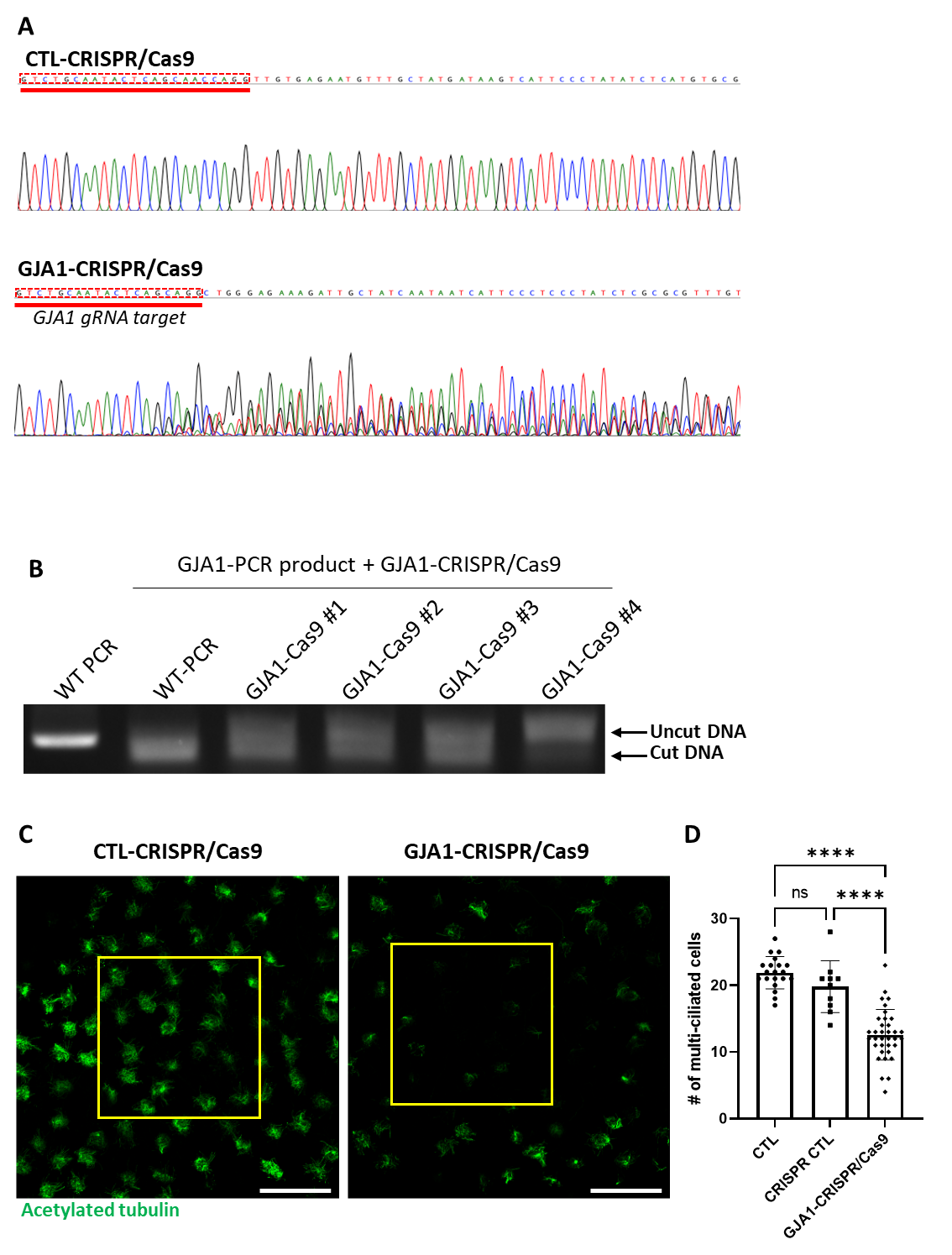

**Figure S4. CRISPR/Cas9-mediated GJA1 knockout shows ciliary defect phenotypes similar to MO-mediated knockdown.**

**A.** The sequencing results of GJA1 genomic DNA PCR products. CRISPR/Cas9-mediated GJA1 deletion mutagenesis effectively disrupted the GJA1 genomic DNA sequences of the GJA1 guide RNA target.

**B.** *In vitro* GJA1-CRISPR/Cas9 reaction with GJA1 PCR products from wild-type or GJA1-CRISPR/Cas9-injected embryos. GJA1-Cas9 failed to cut PCR products from GJA1-Cas9-injected embryos due to mutations at the target site.

**C.** Microinjection of the GJA1-CRISPR/Cas9 disrupted ciliary bundle formation. The ciliary axonemes were labeled with an acetylated tubulin antibody (green). Scale bars: 100 µm.

**D.** Statistical analysis of the number of multiciliated cells per unit area in panel **(C)**. Error bars represent the mean±s.d. P values were determined by one-way ANOVA (P^****^<0.0001, P^ns^>0.05). Raw values are provided in the source data file.

**Supplementary figure 5**

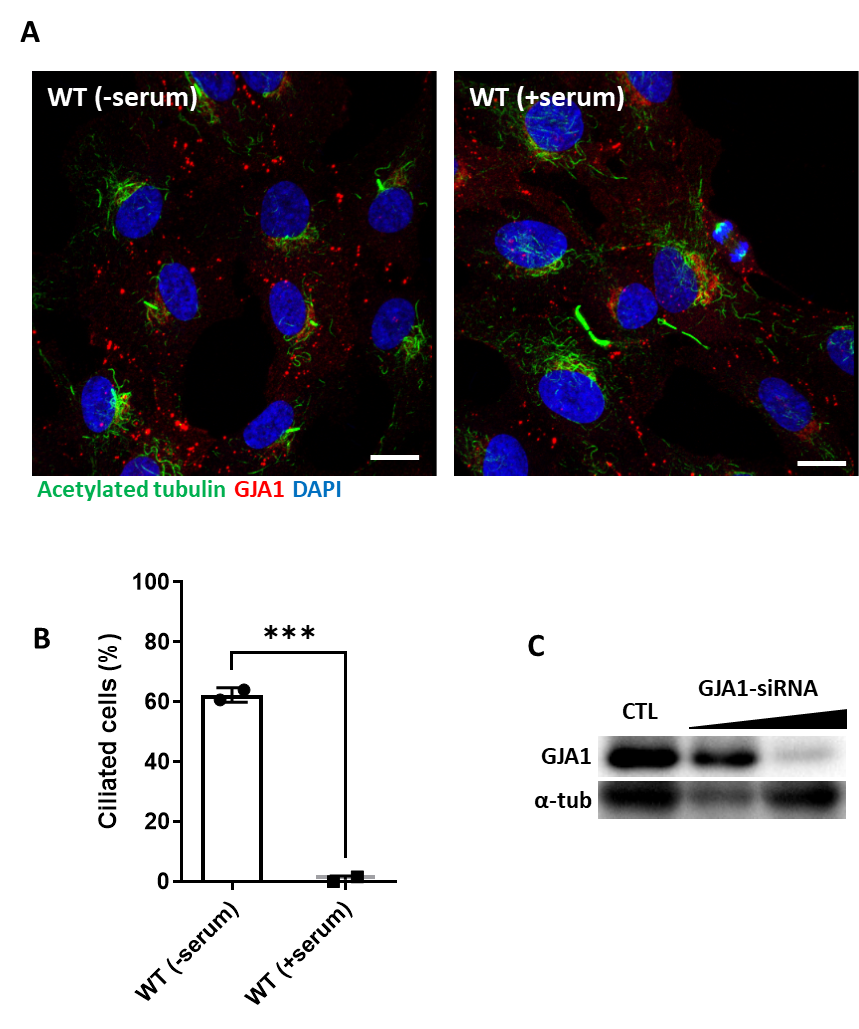

**Figure S5. Ciliation of RPE1 cells by serum starvation and the efficacy of GJA1 siRNA**

**A.** Ciliogenesis of RPE1 was induced by serum starvation. Scale bars: 10 µm.

**B.** The percentage of ciliated cells in panel **(A)** was plotted. Error bars represent the mean±s.d. P values were determined with a two-tailed t-test (P^***^=0.0009). Raw values are provided in the source data file.

**C.** siRNA transfection effectively depleted GJA1 levels in human RPE1 cells. Immunoblotting was performed with an anti-GJA1 antibody and an anti-α-tubulin antibody as a loading control.

**Supplementary figure 6**

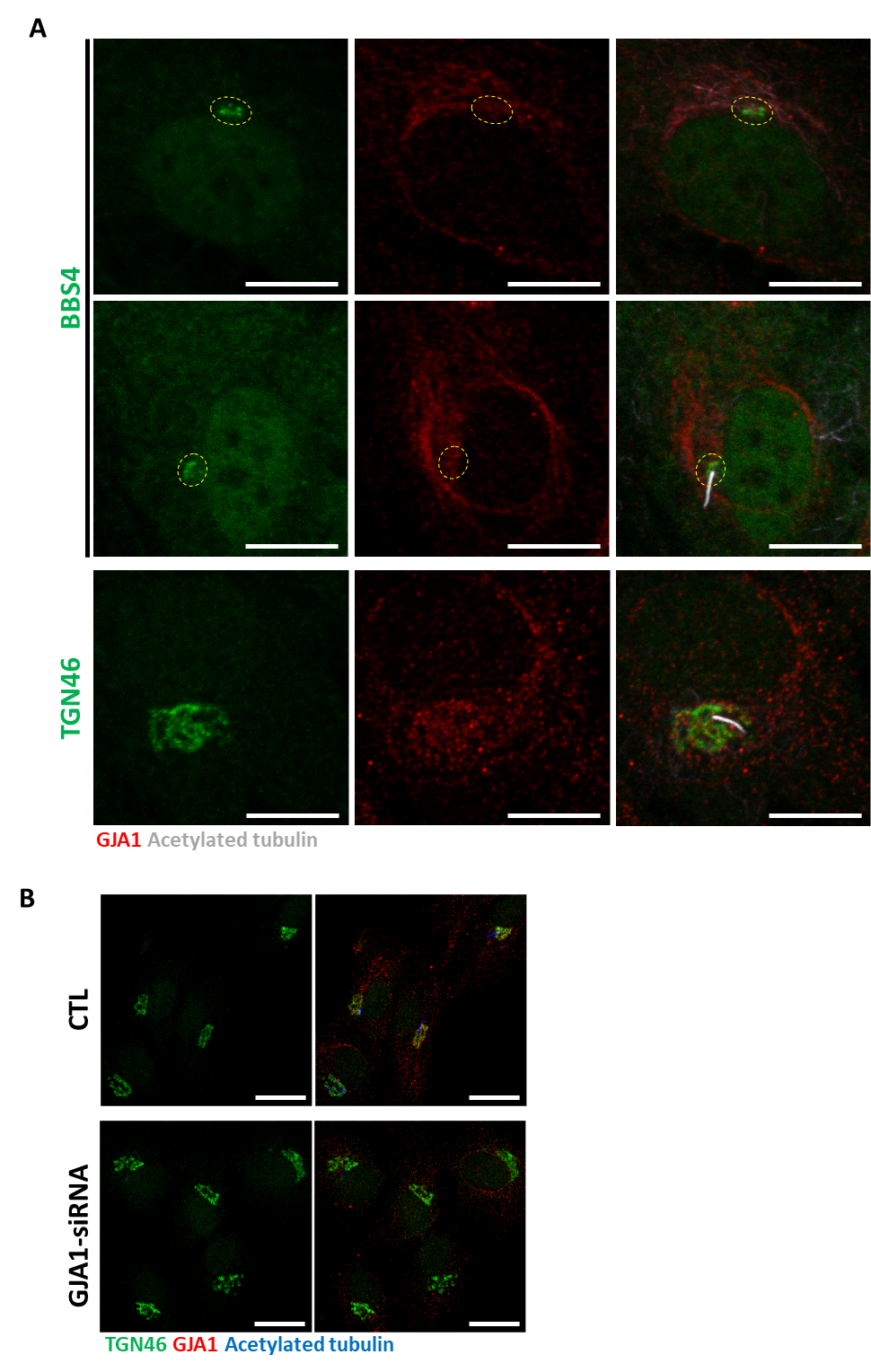

**Figure S6.** **GJA1 localizes to the Golgi complex but GJA1 depletion does not affect Golgi morphology**.

**A.** Serum-starved RPE1 cells were labeled with an anti-BBS4 or anti-TGN46 antibody (green), an anti-GJA1 antibody (red), and an anti-acetylated tubulin antibody (gray). GJA1 localized to the Golgi complex. Scale bars: 10 µm

**B.** RPE1 cells were labeled with an anti-TGN46 antibody (green), an anti-GJA1 antibody (red), and an anti-acetylated tubulin antibody (blue). siRNA-mediated knockdown of GJA1 did not affect the morphology of the Golgi complex. Scale bars: 10 µm.

**Supplementary figure 7**

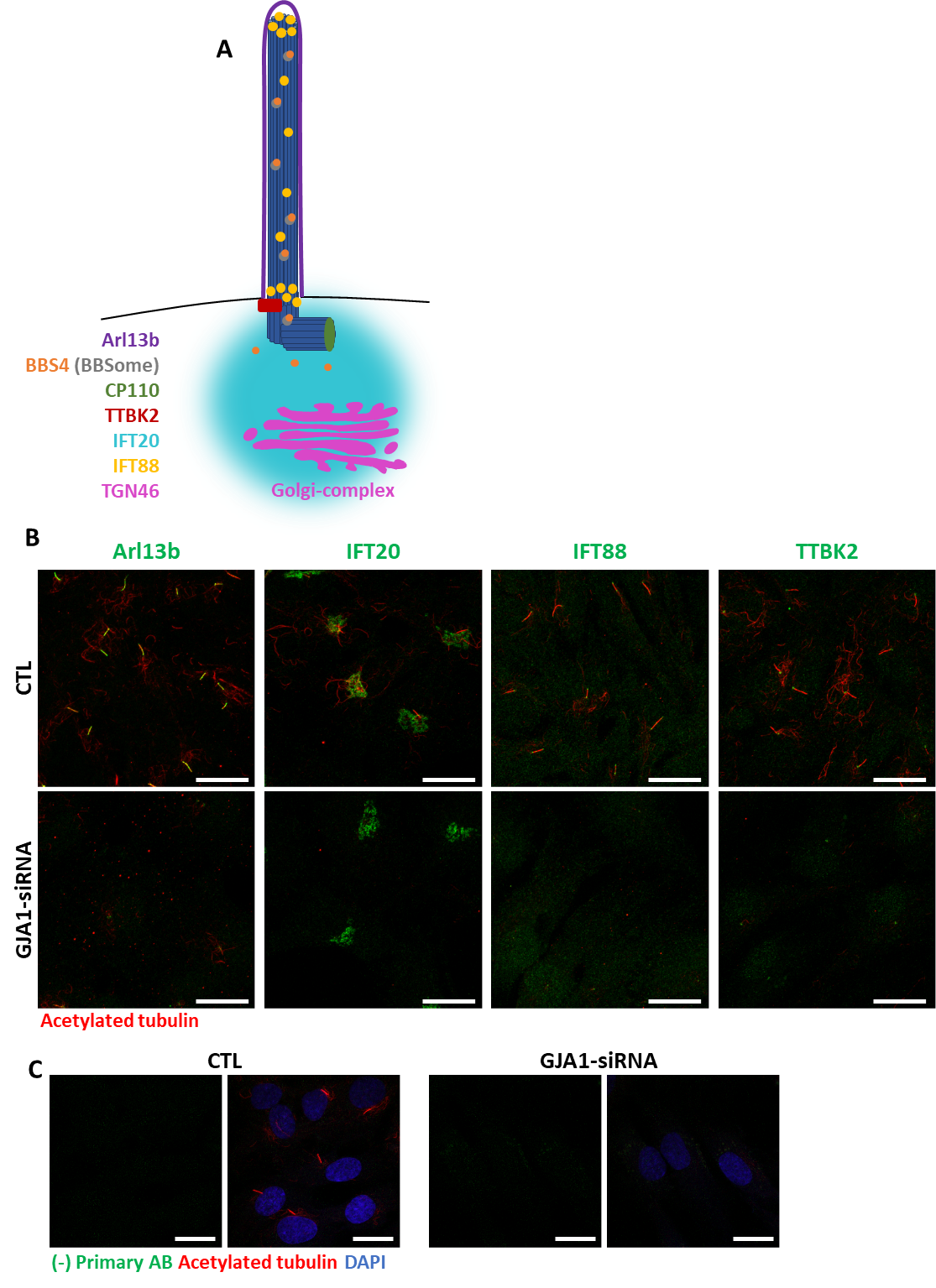

**Figure S7. Subcellular localization of ciliary proteins near the pericentriolar region in GJA1-depleted RPE1 cells.**

**A.** Subcellular localization of ciliary proteins (Arl13B, IFT20, IFT88, and TTBK2) near the pericentriolar region during ciliogenesis.

**B.** RPE1 cells were transfected with GJA1 siRNA and immunostained with acetylated tubulin antibody (red) and anti-Arl13B, anti-IFT20, anti-IFT88, or anti-TTBK2 (green) antibodies. Scale bars: 20 µm.

**C.** RPE1 cells were transfected with GJA1 siRNA and immunostained with an acetylated tubulin antibody (red) and an Alexa-488-conjugated secondary antibody (green) without other primary antibodies as a negative control. Scale bars: 20 µm.

**Supplementary figure 8**

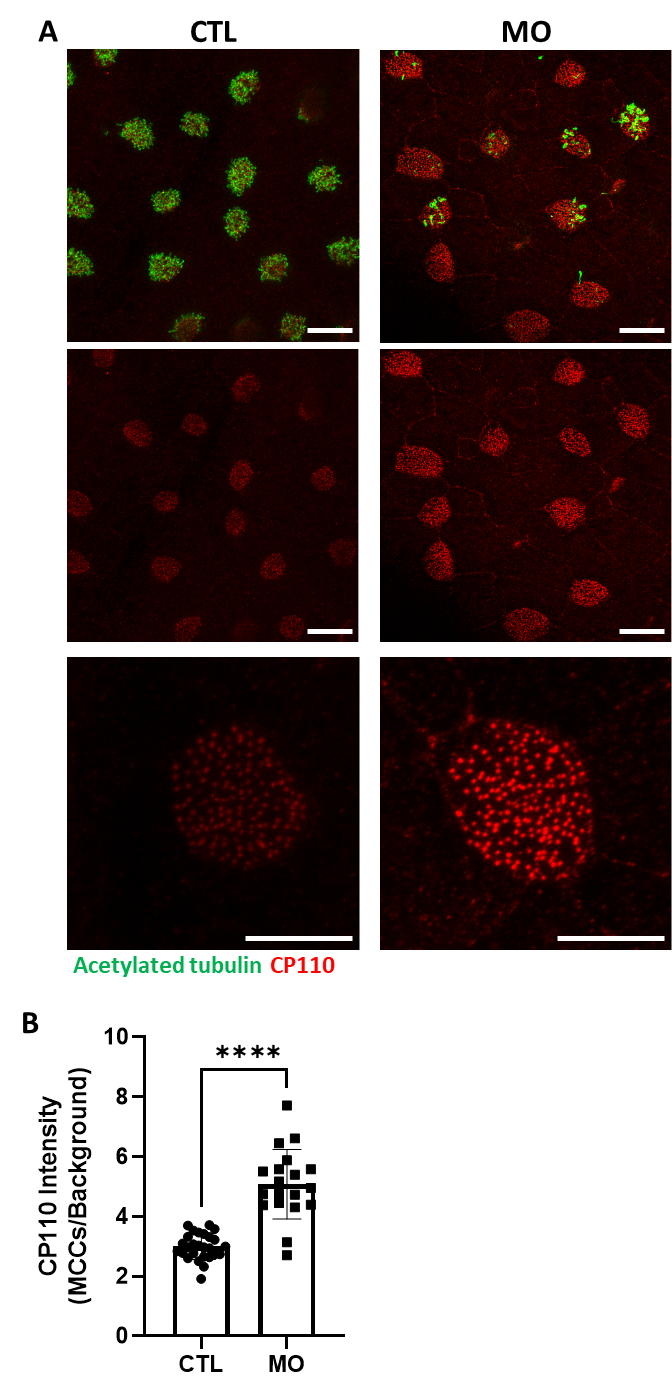

**Figure S8. CP110 localization in multiciliated cells in GJA1-MO-injected *Xenopus* embryos**

**A.** Wild-type and GJA1-MO-injected embryos were stained with CP110 (red) and acetylated tubulin (green) antibodies at stage 30. Scale bars: 20 µm (upper, middle panel), 10 µm (lower panel).

**B.** The relative intensity of CP110 signals in the ciliated cells in panel **(A**) was measured and plotted. Error bars represent the mean±s.d. P values were determined with a two-tailed t-test (P****<0.0001). Raw values are provided in the source data file.

**Supplementary figure 9**

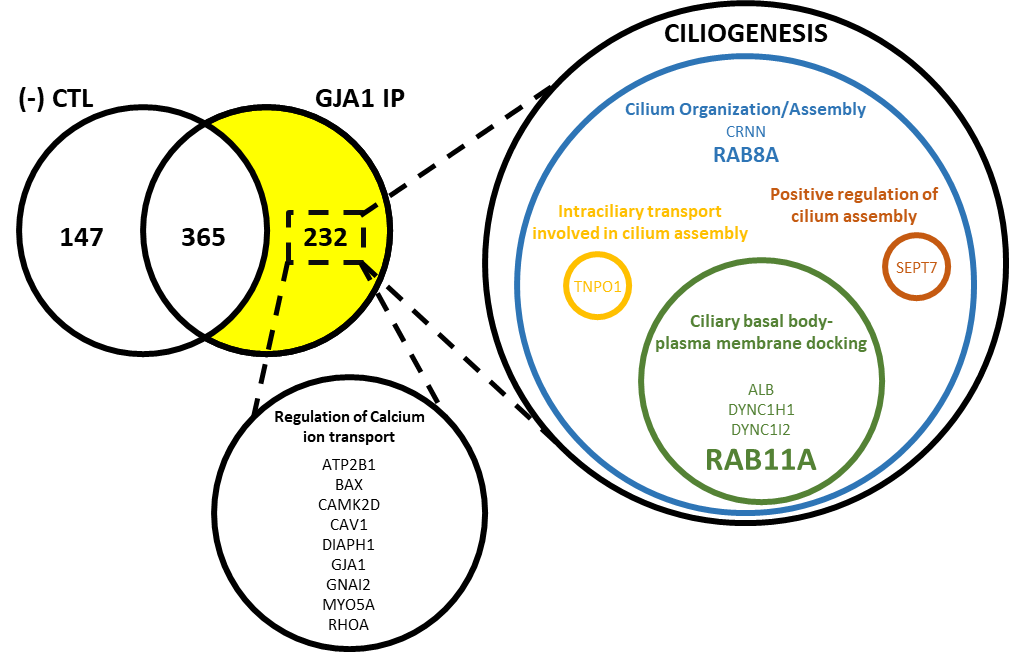

**Figure S9. Identification of GJA1-interacting proteins by IP-MS.**

In total, 232 proteins were detected by immunoprecipitation of GJA1 and mass spectrometry analysis. Gene Ontology term analysis of the proteins identified two distinct groups of proteins involved in Ca^2+^ transport and ciliogenesis.

**Supplementary figure 10**

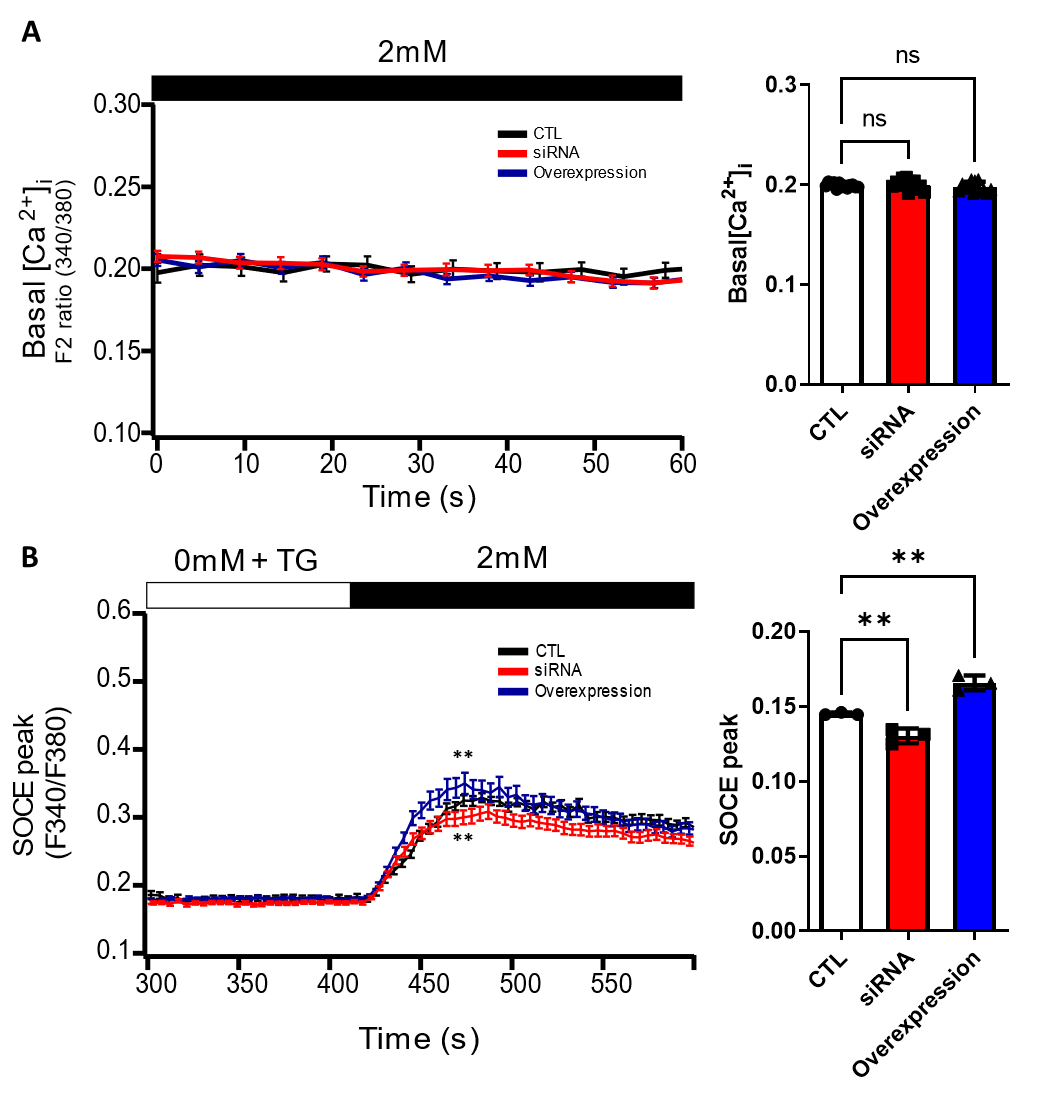

**Figure S10. Basal Ca^2+^ levels and Ca^2+^ influx in GJA1-depleted RPE1 cells.**

**A.** Basal Ca^2+^ levels in RPE1 control (CTL), GJA1 siRNA-transfected, and GJA1 overexpressed cells were monitored by Fura-2 fluorescence ratios. Error bars represent the mean±s.d. P values were determined by one-way ANOVA (P^ns^>0.05). Raw values are provided in the source data file.

**B.** Ca^2+^ influx in RPE1 CTL, GJA1 siRNA-transfected, and GJA1-overexpressed cells was monitored by Fura-2 fluorescence ratios. Error bars represent the mean±s.d. P values were determined by one-way ANOVA (P^**^=0.0076 (siRNA), P^**^=0.0014 (overexpression)). Raw values are provided in the source data file.

**Supplementary figure 11**

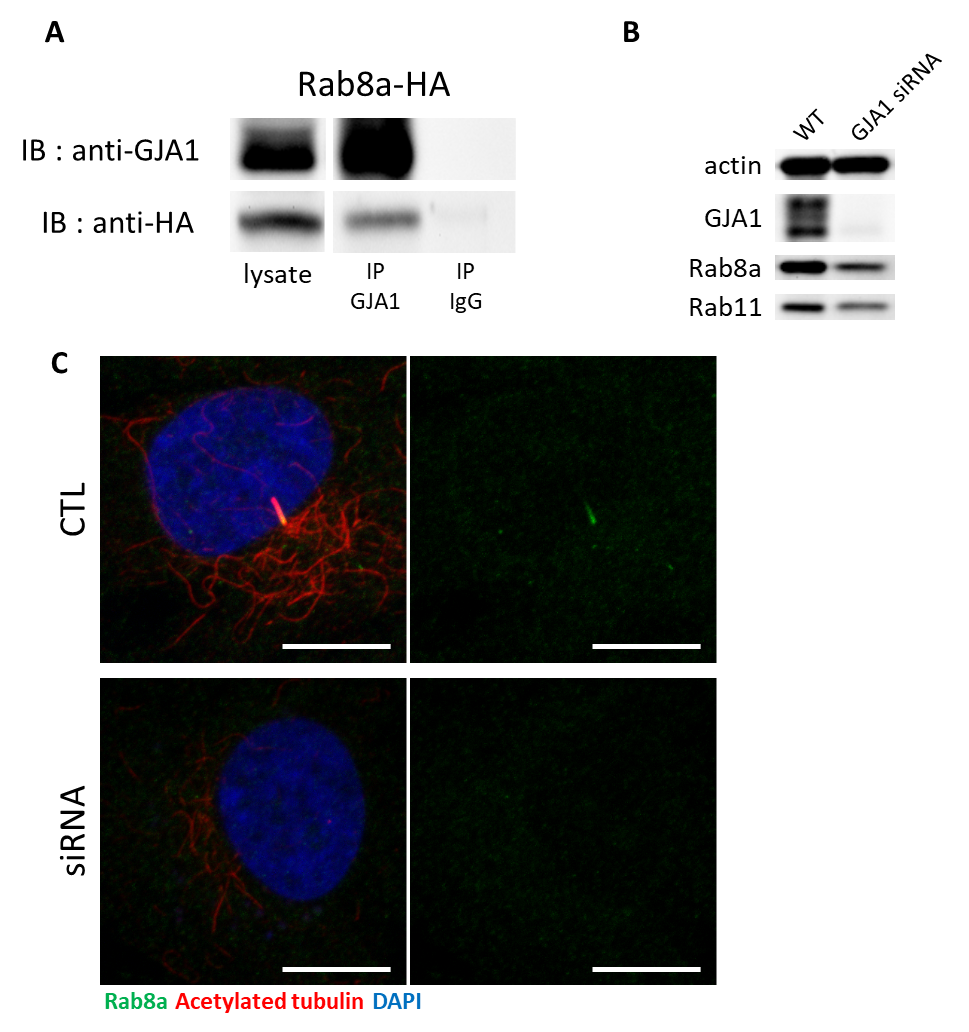

**Figure S11. GJA1 interacts with Rab8a.**

A. HA-tagged Rab8a protein was co-immunoprecipitated with GJA1. Immunoblotting (IB) was performed with anti-GJA1 and anti-HA antibodies.

B. Rab8a, Rab11, and GJA1 expression levels in GJA1 siRNA-transfected RPE1 cells were analyzed by immunoblotting. Actin was used as a loading control.

C. siRNA-mediated knockdown of GJA1 inhibited Rab8a localization in RPE1 cells. RPE1 cells were labeled with an anti-Rab8a antibody (green), an anti-acetylated tubulin antibody (red), and DAPI (blue). Scale bars: 10 µm.

**Supplementary Movi1 1. Heartbeat of wildtype embryos**

**Supplementary Movie 2. Heartbeat of GJA1 morphant embryos**

**Table S1. GJA1 IP-MS results** Ciliogenesis genes (**bold**) and Ca^2+^ transport genes (*italics*) are indicated.

| **Alternate ID** | **Identified information** |
| --- | --- |
| TUBB2A | Tubulin beta-2A chain OS=Homo sapiens GN=TUBB2A PE=1 SV=1 |
| *GJA1* | *Gap junction alpha-1 protein OS=Homo sapiens GN=GJA1 PE=1 SV=2* |
| TRIM21 | E3 ubiquitin-protein ligase TRIM21 OS=Homo sapiens GN=TRIM21 PE=1 SV=1 |
| SPTAN1 | Isoform 2 of Spectrin alpha chain, non-erythrocytic 1 OS=Homo sapiens GN=SPTAN1 |
| MYOF | Myoferlin OS=Homo sapiens GN=MYOF PE=1 SV=1 |
| MPRIP | Isoform 2 of Myosin phosphatase Rho-interacting protein OS=Homo sapiens GN=MPRIP |
| IGF2R | Cation-independent mannose-6-phosphate receptor OS=Homo sapiens GN=IGF2R PE=1 SV=3 |
| MYH13 | Myosin-13 OS=Homo sapiens GN=MYH13 PE=2 SV=2 |
| ELP4 | Isoform 3 of Elongator complex protein 4 OS=Homo sapiens GN=ELP4 |
| RPN1 | Dolichyl-diphosphooligosaccharide--protein glycosyltransferase subunit 1 OS=Homo sapiens GN=RPN1 PE=1 SV=1 |
| ALCAM | CD166 antigen OS=Homo sapiens GN=ALCAM PE=1 SV=2 |
| **ALB** | **Isoform 2 of Serum albumin OS=Homo sapiens GN=ALB** |
| CK | (X98614) cytokeratin [Homo sapiens] |
| **ALB** | **Isoform 3 of Serum albumin OS=Homo sapiens GN=ALB** |
| ATP1A1 | Sodium/potassium-transporting ATPase subunit alpha-1 OS=Homo sapiens GN=ATP1A1 PE=1 SV=1 |
| ITGA3 | Integrin alpha-3 OS=Homo sapiens GN=ITGA3 PE=1 SV=5 |
| ITGAV | Integrin alpha-V OS=Homo sapiens GN=ITGAV PE=1 SV=2 |
| NPM1 | Isoform 2 of Nucleophosmin OS=Homo sapiens GN=NPM1 |
| KRT15 | Isoform 2 of Keratin, type I cytoskeletal 15 OS=Homo sapiens GN=KRT15 |
| ITGB5 | Integrin beta-5 OS=Homo sapiens GN=ITGB5 PE=1 SV=1 |
| RAB1B | Ras-related protein Rab-1B OS=Homo sapiens GN=RAB1B PE=1 SV=1 |
| SURF4 | Surfeit locus protein 4 OS=Homo sapiens GN=SURF4 PE=1 SV=3 |
| PPP1R18 | Phostensin OS=Homo sapiens GN=PPP1R18 PE=1 SV=1 |
| RPN2 | Dolichyl-diphosphooligosaccharide--protein glycosyltransferase subunit 2 OS=Homo sapiens GN=RPN2 PE=1 SV=3 |
| ELP5 | Elongator complex protein 5 OS=Homo sapiens GN=ELP5 PE=1 SV=2 |
| ELP6 | Elongator complex protein 6 OS=Homo sapiens GN=ELP6 PE=1 SV=1 |
| ARL6IP5 | PRA1 family protein 3 OS=Homo sapiens GN=ARL6IP5 PE=1 SV=1 |
| **DYNC1I2** | **Isoform 2B of Cytoplasmic dynein 1 intermediate chain 2 OS=Homo sapiens GN=DYNC1I2** |
| TMED9 | Transmembrane emp24 domain-containing protein 9 OS=Homo sapiens GN=TMED9 PE=1 SV=2 |
| SCFD1 | Sec1 family domain-containing protein 1 OS=Homo sapiens GN=SCFD1 PE=1 SV=4 |
| ATL3 | Atlastin-3 OS=Homo sapiens GN=ATL3 PE=1 SV=1 |
| XPO1 | Exportin-1 OS=Homo sapiens GN=XPO1 PE=1 SV=1 |
| TFRC | Transferrin receptor protein 1 OS=Homo sapiens GN=TFRC PE=1 SV=2 |
| TXLNA | Alpha-taxilin OS=Homo sapiens GN=TXLNA PE=1 SV=3 |
| RAB7A | Ras-related protein Rab-7a OS=Homo sapiens GN=RAB7A PE=1 SV=1 |
| DDOST | Dolichyl-diphosphooligosaccharide--protein glycosyltransferase 48 kDa subunit OS=Homo sapiens GN=DDOST PE=1 SV=4 |
| *CAMK2D* | *Isoform Delta 3 of Calcium/calmodulin-dependent protein kinase type II subunit delta OS=Homo sapiens GN=CAMK2D* |
| *RHOA* | *Transforming protein RhoA OS=Homo sapiens GN=RHOA PE=1 SV=1* |
| CD44 | CD44 antigen OS=Homo sapiens GN=CD44 PE=1 SV=3 |
| LGALS3BP | Galectin-3-binding protein OS=Homo sapiens GN=LGALS3BP PE=1 SV=1 |
| MRC2 | C-type mannose receptor 2 OS=Homo sapiens GN=MRC2 PE=1 SV=2 |
| CNN2 | Calponin-2 OS=Homo sapiens GN=CNN2 PE=1 SV=4 |
| **RAB8A** | **Ras-related protein Rab-8A OS=Homo sapiens GN=RAB8A PE=1 SV=1** |
| RTN3 | Reticulon-3 OS=Homo sapiens GN=RTN3 PE=1 SV=2 |
| **TNPO1** | **Isoform 3 of Transportin-1 OS=Homo sapiens GN=TNPO1** |
| GCN1 | eIF-2-alpha kinase activator GCN1 OS=Homo sapiens GN=GCN1 PE=1 SV=6 |
| GNS | N-acetylglucosamine-6-sulfatase OS=Homo sapiens GN=GNS PE=1 SV=3 |
| SDHA | Succinate dehydrogenase [ubiquinone] flavoprotein subunit, mitochondrial OS=Homo sapiens GN=SDHA PE=1 SV=2 |
| *ATP2B1* | *Plasma membrane calcium-transporting ATPase 1 OS=Homo sapiens GN=ATP2B1 PE=1 SV=3* |
| TMED4 | Transmembrane emp24 domain-containing protein 4 OS=Homo sapiens GN=TMED4 PE=1 SV=1 |
| SFPQ | Splicing factor, proline- and glutamine-rich OS=Homo sapiens GN=SFPQ PE=1 SV=2 |
| LMAN2 | Vesicular integral-membrane protein VIP36 OS=Homo sapiens GN=LMAN2 PE=1 SV=1 |
| LRRFIP2 | Isoform 4 of Leucine-rich repeat flightless-interacting protein 2 OS=Homo sapiens GN=LRRFIP2 |
| PPT1 | Palmitoyl-protein thioesterase 1 OS=Homo sapiens GN=PPT1 PE=1 SV=1 |
| FUBP1 | Far upstream element-binding protein 1 OS=Homo sapiens GN=FUBP1 PE=1 SV=3 |
| **RAB11A** | **Ras-related protein Rab-11A OS=Homo sapiens GN=RAB11A PE=1 SV=3** |
| GLG1 | Golgi apparatus protein 1 OS=Homo sapiens GN=GLG1 PE=1 SV=2 |
| ESYT1 | Isoform 2 of Extended synaptotagmin-1 OS=Homo sapiens GN=ESYT1 |
| LMNB1 | Lamin-B1 OS=Homo sapiens GN=LMNB1 PE=1 SV=2 |
| TMED10 | Transmembrane emp24 domain-containing protein 10 OS=Homo sapiens GN=TMED10 PE=1 SV=2 |
| *BAX* | *Isoform Sigma of Apoptosis regulator BAX OS=Homo sapiens GN=BAX* |
| SCCPDH | Saccharopine dehydrogenase-like oxidoreductase OS=Homo sapiens GN=SCCPDH PE=1 SV=1 |
| CD81 | CD81 antigen OS=Homo sapiens GN=CD81 PE=1 SV=1 |
| ITGA5 | Integrin alpha-5 OS=Homo sapiens GN=ITGA5 PE=1 SV=2 |
| RAB5A | Ras-related protein Rab-5A OS=Homo sapiens GN=RAB5A PE=1 SV=2 |
| IGLL5 | Immunoglobulin lambda-like polypeptide 5 OS=Homo sapiens GN=IGLL5 PE=2 SV=2 |
| COL27A1 | Collagen alpha-1(XXVII) chain OS=Homo sapiens GN=COL27A1 PE=1 SV=1 |
| PLBD2 | Putative phospholipase B-like 2 OS=Homo sapiens GN=PLBD2 PE=1 SV=2 |
| EPHA2 | Ephrin type-A receptor 2 OS=Homo sapiens GN=EPHA2 PE=1 SV=2 |
| MAP1B | Microtubule-associated protein 1B OS=Homo sapiens GN=MAP1B PE=1 SV=2 |
| UQCRC2 | Cytochrome b-c1 complex subunit 2, mitochondrial OS=Homo sapiens GN=UQCRC2 PE=1 SV=3 |
| SRP68 | Signal recognition particle subunit SRP68 OS=Homo sapiens GN=SRP68 PE=1 SV=2 |
| GSR | Isoform 4 of Glutathione reductase, mitochondrial OS=Homo sapiens GN=GSR |
| TGOLN2 | Isoform 5 of Trans-Golgi network integral membrane protein 2 OS=Homo sapiens GN=TGOLN2 |
| GIPC1 | PDZ domain-containing protein GIPC1 OS=Homo sapiens GN=GIPC1 PE=1 SV=2 |
| SERPINB3 | Serpin B3 OS=Homo sapiens GN=SERPINB3 PE=1 SV=2 |
| DBNL | Isoform 2 of Drebrin-like protein OS=Homo sapiens GN=DBNL |
| PCYOX1 | Isoform 2 of Prenylcysteine oxidase 1 OS=Homo sapiens GN=PCYOX1 |
| TICAM2 | Isoform 2 of TIR domain-containing adapter molecule 2 OS=Homo sapiens GN=TICAM2 |
| PIGR | Polymeric immunoglobulin receptor OS=Homo sapiens GN=PIGR PE=1 SV=4 |
| ATP5F1 | ATP synthase F(0) complex subunit B1, mitochondrial OS=Homo sapiens GN=ATP5F1 PE=1 SV=2 |
| HACD3 | Very-long-chain (3R)-3-hydroxyacyl-CoA dehydratase 3 OS=Homo sapiens GN=HACD3 PE=1 SV=2 |
| SCARB2 | Isoform 2 of Lysosome membrane protein 2 OS=Homo sapiens GN=SCARB2 |
| PSMA4 | Proteasome subunit alpha type-4 OS=Homo sapiens GN=PSMA4 PE=1 SV=1 |
| SLC25A3 | Isoform B of Phosphate carrier protein, mitochondrial OS=Homo sapiens GN=SLC25A3 |
| HLA-A | HLA class I histocompatibility antigen, A-2 alpha chain OS=Homo sapiens GN=HLA-A PE=1 SV=1 |
| SERPINB12 | Serpin B12 OS=Homo sapiens GN=SERPINB12 PE=1 SV=1 |
| TMEM106B | Transmembrane protein 106B OS=Homo sapiens GN=TMEM106B PE=1 SV=2 |
| RAB2A | Ras-related protein Rab-2A OS=Homo sapiens GN=RAB2A PE=1 SV=1 |
| AHCY | Adenosylhomocysteinase OS=Homo sapiens GN=AHCY PE=1 SV=4 |
| PDLIM2 | Isoform 5 of PDZ and LIM domain protein 2 OS=Homo sapiens GN=PDLIM2 |
| S100A8 | Protein S100-A8 OS=Homo sapiens GN=S100A8 PE=1 SV=1 |
| EIF3F | Eukaryotic translation initiation factor 3 subunit F OS=Homo sapiens GN=EIF3F PE=1 SV=1 |
| TBC1D15 | Isoform 3 of TBC1 domain family member 15 OS=Homo sapiens GN=TBC1D15 |
| *CAV1* | *Caveolin-1 OS=Homo sapiens GN=CAV1 PE=1 SV=4* |
| SCAMP1 | Secretory carrier-associated membrane protein 1 OS=Homo sapiens GN=SCAMP1 PE=1 SV=2 |
| HYOU1 | Hypoxia up-regulated protein 1 OS=Homo sapiens GN=HYOU1 PE=1 SV=1 |
| HTRA1 | Serine protease HTRA1 OS=Homo sapiens GN=HTRA1 PE=1 SV=1 |
| RPS12 | 40S ribosomal protein S12 OS=Homo sapiens GN=RPS12 PE=1 SV=3 |
| PSMC5 | 26S protease regulatory subunit 8 OS=Homo sapiens GN=PSMC5 PE=1 SV=1 |
| TPD52L2 | Tumor protein D54 OS=Homo sapiens GN=TPD52L2 PE=1 SV=2 |
| RDH11 | Retinol dehydrogenase 11 OS=Homo sapiens GN=RDH11 PE=1 SV=2 |
| TM9SF3 | Transmembrane 9 superfamily member 3 OS=Homo sapiens GN=TM9SF3 PE=1 SV=2 |
| PDLIM4 | PDZ and LIM domain protein 4 OS=Homo sapiens GN=PDLIM4 PE=1 SV=2 |
| GOLT1B | Vesicle transport protein GOT1B OS=Homo sapiens GN=GOLT1B PE=1 SV=1 |
| CSN3 | KAPPA CASEIN PRECURSOR. - BOS TAURUS (BOVINE). |
| RER1 | Protein RER1 OS=Homo sapiens GN=RER1 PE=1 SV=1 |
| ATP6V1E1 | Isoform 2 of V-type proton ATPase subunit E 1 OS=Homo sapiens GN=ATP6V1E1 |
| ADAM9 | Disintegrin and metalloproteinase domain-containing protein 9 OS=Homo sapiens GN=ADAM9 PE=1 SV=1 |
| STMN1 | Stathmin OS=Homo sapiens GN=STMN1 PE=1 SV=3 |
| CD63 | Isoform 3 of CD63 antigen OS=Homo sapiens GN=CD63 |
| *MYO5A* | *Unconventional myosin-Va OS=Homo sapiens GN=MYO5A PE=1 SV=2* |
| RPL31 | 60S ribosomal protein L31 OS=Homo sapiens GN=RPL31 PE=1 SV=1 |
| B2M | Beta-2-microglobulin OS=Homo sapiens GN=B2M PE=1 SV=1 |
| **DYNC1H1** | **Cytoplasmic dynein 1 heavy chain 1 OS=Homo sapiens GN=DYNC1H1 PE=1 SV=5** |
| ERAP1 | Endoplasmic reticulum aminopeptidase 1 OS=Homo sapiens GN=ERAP1 PE=1 SV=3 |
| GBF1 | Golgi-specific brefeldin A-resistance guanine nucleotide exchange factor 1 OS=Homo sapiens GN=GBF1 PE=1 SV=2 |
| NUP153 | Nuclear pore complex protein Nup153 OS=Homo sapiens GN=NUP153 PE=1 SV=2 |
| COL3A1 | Collagen alpha-1(III) chain OS=Homo sapiens GN=COL3A1 PE=1 SV=4 |
| PSMD4 | 26S proteasome non-ATPase regulatory subunit 4 OS=Homo sapiens GN=PSMD4 PE=1 SV=1 |
| LPCAT2 | Lysophosphatidylcholine acyltransferase 2 OS=Homo sapiens GN=LPCAT2 PE=1 SV=1 |
| PSMD11 | 26S proteasome non-ATPase regulatory subunit 11 OS=Homo sapiens GN=PSMD11 PE=1 SV=3 |
| PSMC3 | 26S protease regulatory subunit 6A OS=Homo sapiens GN=PSMC3 PE=1 SV=3 |
| USP5 | Ubiquitin carboxyl-terminal hydrolase 5 OS=Homo sapiens GN=USP5 PE=1 SV=2 |
| PNP | Purine nucleoside phosphorylase OS=Homo sapiens GN=PNP PE=1 SV=2 |
| USP31 | Ubiquitin carboxyl-terminal hydrolase 31 OS=Homo sapiens GN=USP31 PE=2 SV=2 |
| ATP6V1C1 | V-type proton ATPase subunit C 1 OS=Homo sapiens GN=ATP6V1C1 PE=1 SV=4 |
| MAN2B2 | Epididymis-specific alpha-mannosidase OS=Homo sapiens GN=MAN2B2 PE=1 SV=4 |
| SLC3A2 | 4F2 cell-surface antigen heavy chain OS=Homo sapiens GN=SLC3A2 PE=1 SV=3 |
| RPS10 | 40S ribosomal protein S10 OS=Homo sapiens GN=RPS10 PE=1 SV=1 |
| ACSL3 | Long-chain-fatty-acid--CoA ligase 3 OS=Homo sapiens GN=ACSL3 PE=1 SV=3 |
| RETSAT | All-trans-retinol 13,14-reductase OS=Homo sapiens GN=RETSAT PE=1 SV=2 |
| SENP5 | Sentrin-specific protease 5 OS=Homo sapiens GN=SENP5 PE=1 SV=3 |
| TRAF3 | TNF receptor-associated factor 3 OS=Homo sapiens GN=TRAF3 PE=1 SV=2 |
| ATP2A2 | Sarcoplasmic/endoplasmic reticulum calcium ATPase 2 OS=Homo sapiens GN=ATP2A2 PE=1 SV=1 |
| **SEPT7** | **Septin-7 OS=Homo sapiens GN=SEPT7 PE=1 SV=2** |
| HNRNPM | Heterogeneous nuclear ribonucleoprotein M OS=Homo sapiens GN=HNRNPM PE=1 SV=3 |
| ALDH3A2 | Isoform 2 of Fatty aldehyde dehydrogenase OS=Homo sapiens GN=ALDH3A2 |
| DYNC1LI1 | Cytoplasmic dynein 1 light intermediate chain 1 OS=Homo sapiens GN=DYNC1LI1 PE=1 SV=3 |
| CSN1S2 | ALPHA-S2 CASEIN PRECURSOR. - BOS TAURUS (BOVINE). |
| MYADM | Myeloid-associated differentiation marker OS=Homo sapiens GN=MYADM PE=1 SV=2 |
| MOGS | Isoform 2 of Mannosyl-oligosaccharide glucosidase OS=Homo sapiens GN=MOGS |
| BCAP31 | B-cell receptor-associated protein 31 OS=Homo sapiens GN=BCAP31 PE=1 SV=3 |
| COMT | Catechol O-methyltransferase OS=Homo sapiens GN=COMT PE=1 SV=2 |
| SEC31A | Protein transport protein Sec31A OS=Homo sapiens GN=SEC31A PE=1 SV=3 |
| ADRM1 | Proteasomal ubiquitin receptor ADRM1 OS=Homo sapiens GN=ADRM1 PE=1 SV=2 |
| KRT9 | ct\|1082558-DECOY\|S41161 keratin 9, cytoskeletal - human |
| EEA1 | Early endosome antigen 1 OS=Homo sapiens GN=EEA1 PE=1 SV=2 |
| SLC2A1 | Solute carrier family 2, facilitated glucose transporter member 1 OS=Homo sapiens GN=SLC2A1 PE=1 SV=2 |
| HSPH1 | Heat shock protein 105 kDa OS=Homo sapiens GN=HSPH1 PE=1 SV=1 |
| PIP | Prolactin-inducible protein OS=Homo sapiens GN=PIP PE=1 SV=1 |
| MFGE8 | Lactadherin OS=Homo sapiens GN=MFGE8 PE=1 SV=2 |
| FASN | Fatty acid synthase OS=Homo sapiens GN=FASN PE=1 SV=3 |
| ANKS1A | Ankyrin repeat and SAM domain-containing protein 1A OS=Homo sapiens GN=ANKS1A PE=1 SV=4 |
| LMAN1 | Protein ERGIC-53 OS=Homo sapiens GN=LMAN1 PE=1 SV=2 |
| VDAC1 | Voltage-dependent anion-selective channel protein 1 OS=Homo sapiens GN=VDAC1 PE=1 SV=2 |
| MLEC | Malectin OS=Homo sapiens GN=MLEC PE=1 SV=1 |
| SEL1L | Protein sel-1 homolog 1 OS=Homo sapiens GN=SEL1L PE=1 SV=3 |
| ADAM10 | Disintegrin and metalloproteinase domain-containing protein 10 OS=Homo sapiens GN=ADAM10 PE=1 SV=1 |
| *DIAPH1* | *Isoform 3 of Protein diaphanous homolog 1 OS=Homo sapiens GN=DIAPH1* |
| ESYT2 | Isoform 2 of Extended synaptotagmin-2 OS=Homo sapiens GN=ESYT2 |
| NNT | NAD(P) transhydrogenase, mitochondrial OS=Homo sapiens GN=NNT PE=1 SV=3 |
| GAS6 | Growth arrest-specific protein 6 OS=Homo sapiens GN=GAS6 PE=1 SV=2 |
| RAC1 | Isoform B of Ras-related C3 botulinum toxin substrate 1 OS=Homo sapiens GN=RAC1 |
| RPL34 | 60S ribosomal protein L34 OS=Homo sapiens GN=RPL34 PE=1 SV=3 |
| CSTB | Cystatin-B OS=Homo sapiens GN=CSTB PE=1 SV=2 |
| CTSZ | Cathepsin Z OS=Homo sapiens GN=CTSZ PE=1 SV=1 |
| TMEM43 | Transmembrane protein 43 OS=Homo sapiens GN=TMEM43 PE=1 SV=1 |
| SH3BP4 | SH3 domain-binding protein 4 OS=Homo sapiens GN=SH3BP4 PE=1 SV=1 |
| CBR1 | Carbonyl reductase [NADPH] 1 OS=Homo sapiens GN=CBR1 PE=1 SV=3 |
| FAP | Prolyl endopeptidase FAP OS=Homo sapiens GN=FAP PE=1 SV=5 |
| NEDD4L | E3 ubiquitin-protein ligase NEDD4-like OS=Homo sapiens GN=NEDD4L PE=1 SV=2 |
| HIP1 | Huntingtin-interacting protein 1 OS=Homo sapiens GN=HIP1 PE=1 SV=5 |
| GPD2 | Glycerol-3-phosphate dehydrogenase, mitochondrial OS=Homo sapiens GN=GPD2 PE=1 SV=3 |
| HADHB | Trifunctional enzyme subunit beta, mitochondrial OS=Homo sapiens GN=HADHB PE=1 SV=3 |
| PSMB6 | Proteasome subunit beta type-6 OS=Homo sapiens GN=PSMB6 PE=1 SV=4 |
| STX7 | Syntaxin-7 OS=Homo sapiens GN=STX7 PE=1 SV=4 |
| ITGA2 | Integrin alpha-2 OS=Homo sapiens GN=ITGA2 PE=1 SV=1 |
| PDLIM1 | PDZ and LIM domain protein 1 OS=Homo sapiens GN=PDLIM1 PE=1 SV=4 |
| GOLIM4 | Golgi integral membrane protein 4 OS=Homo sapiens GN=GOLIM4 PE=1 SV=1 |
| ST13 | Hsc70-interacting protein OS=Homo sapiens GN=ST13 PE=1 SV=2 |
| GPR180 | Integral membrane protein GPR180 OS=Homo sapiens GN=GPR180 PE=2 SV=1 |
| IPO7 | Importin-7 OS=Homo sapiens GN=IPO7 PE=1 SV=1 |
| FARSB | Phenylalanine--tRNA ligase beta subunit OS=Homo sapiens GN=FARSB PE=1 SV=3 |
| IPO5 | Isoform 3 of Importin-5 OS=Homo sapiens GN=IPO5 |
| IMPAD1 | Inositol monophosphatase 3 OS=Homo sapiens GN=IMPAD1 PE=1 SV=1 |
| ETFB | Electron transfer flavoprotein subunit beta OS=Homo sapiens GN=ETFB PE=1 SV=3 |
| BLMH | Bleomycin hydrolase OS=Homo sapiens GN=BLMH PE=1 SV=1 |
| S100A14 | Protein S100-A14 OS=Homo sapiens GN=S100A14 PE=1 SV=1 |
| TOMM70 | Mitochondrial import receptor subunit TOM70 OS=Homo sapiens GN=TOMM70 PE=1 SV=1 |
| GLIPR1L1 | sp\|Q6UWM5-DECOY\|GPRL1_HUMAN GLIPR1-like protein 1 OS=Homo sapiens GN=GLIPR1L1 PE=1... |
| DDX60 | Probable ATP-dependent RNA helicase DDX60 OS=Homo sapiens GN=DDX60 PE=1 SV=3 |
| *GNAI2* | *Guanine nucleotide-binding protein G(i) subunit alpha-2 OS=Homo sapiens GN=GNAI2 PE=1 SV=3* |
| SLC25A6 | ADP/ATP translocase 3 OS=Homo sapiens GN=SLC25A6 PE=1 SV=4 |
| FTL | Ferritin light chain OS=Homo sapiens GN=FTL PE=1 SV=2 |
| JMY | Isoform 2 of Junction-mediating and -regulatory protein OS=Homo sapiens GN=JMY |
| ESD | S-formylglutathione hydrolase OS=Homo sapiens GN=ESD PE=1 SV=2 |
| FAM3C | Protein FAM3C OS=Homo sapiens GN=FAM3C PE=1 SV=1 |
| PSAP | Prosaposin OS=Homo sapiens GN=PSAP PE=1 SV=2 |
| CYB5R3 | NADH-cytochrome b5 reductase 3 OS=Homo sapiens GN=CYB5R3 PE=1 SV=3 |
| EIF4A2 | Eukaryotic initiation factor 4A-II OS=Homo sapiens GN=EIF4A2 PE=1 SV=2 |
| CD276 | CD276 antigen OS=Homo sapiens GN=CD276 PE=1 SV=1 |
| **CRNN** | **Cornulin OS=Homo sapiens GN=CRNN PE=1 SV=1** |
| HLA-A | HLA class I histocompatibility antigen, A-1 alpha chain OS=Homo sapiens GN=HLA-A PE=1 SV=1 |
| ERLIN1 | Erlin-1 OS=Homo sapiens GN=ERLIN1 PE=1 SV=1 |
| SEC11A | Signal peptidase complex catalytic subunit SEC11A OS=Homo sapiens GN=SEC11A PE=1 SV=1 |
| ERGIC3 | Endoplasmic reticulum-Golgi intermediate compartment protein 3 OS=Homo sapiens GN=ERGIC3 PE=1 SV=1 |
| MARS | Methionine--tRNA ligase, cytoplasmic OS=Homo sapiens GN=MARS PE=1 SV=2 |
| ACAT1 | Acetyl-CoA acetyltransferase, mitochondrial OS=Homo sapiens GN=ACAT1 PE=1 SV=1 |
| PSMD2 | 26S proteasome non-ATPase regulatory subunit 2 OS=Homo sapiens GN=PSMD2 PE=1 SV=3 |
| HNRNPCL2 | Heterogeneous nuclear ribonucleoprotein C-like 2 OS=Homo sapiens GN=HNRNPCL2 PE=1 SV=1 |
| MAN2B1 | Lysosomal alpha-mannosidase OS=Homo sapiens GN=MAN2B1 PE=1 SV=3 |
| EIF3H | Eukaryotic translation initiation factor 3 subunit H OS=Homo sapiens GN=EIF3H PE=1 SV=1 |
| DNAJB6 | Isoform B of DnaJ homolog subfamily B member 6 OS=Homo sapiens GN=DNAJB6 |
| IGKC | Ig kappa chain C region OS=Homo sapiens GN=IGKC PE=1 SV=1 |
| LTF | Lactotransferrin OS=Homo sapiens GN=LTF PE=1 SV=6 |
| SMPD1 | Sphingomyelin phosphodiesterase OS=Homo sapiens GN=SMPD1 PE=1 SV=4 |
| PSMB1 | Proteasome subunit beta type-1 OS=Homo sapiens GN=PSMB1 PE=1 SV=2 |
| PSMA3 | Isoform 2 of Proteasome subunit alpha type-3 OS=Homo sapiens GN=PSMA3 |
| CALML3 | Calmodulin-like protein 3 OS=Homo sapiens GN=CALML3 PE=1 SV=2 |
| MAT2A | S-adenosylmethionine synthase isoform type-2 OS=Homo sapiens GN=MAT2A PE=1 SV=1 |
| RPS5 | 40S ribosomal protein S5 OS=Homo sapiens GN=RPS5 PE=1 SV=4 |
| TMEM33 | Transmembrane protein 33 OS=Homo sapiens GN=TMEM33 PE=1 SV=2 |
| LBR | Lamin-B receptor OS=Homo sapiens GN=LBR PE=1 SV=2 |
| SPCS2 | Signal peptidase complex subunit 2 OS=Homo sapiens GN=SPCS2 PE=1 SV=3 |
| SF3B4 | Splicing factor 3B subunit 4 OS=Homo sapiens GN=SF3B4 PE=1 SV=1 |
| SLC30A7 | Zinc transporter 7 OS=Homo sapiens GN=SLC30A7 PE=2 SV=1 |
| SRPK1 | SRSF protein kinase 1 OS=Homo sapiens GN=SRPK1 PE=1 SV=2 |
| SDF2 | Stromal cell-derived factor 2 OS=Homo sapiens GN=SDF2 PE=1 SV=2 |
| VKORC1 | Vitamin K epoxide reductase complex subunit 1 OS=Homo sapiens GN=VKORC1 PE=1 SV=1 |
| TMEM109 | Transmembrane protein 109 OS=Homo sapiens GN=TMEM109 PE=1 SV=1 |
| NSFL1C | Isoform 2 of NSFL1 cofactor p47 OS=Homo sapiens GN=NSFL1C |
